# Discovering Heterogeneous Neurodegenerative Disease Patterns From MRI Data for Improved Prediction

**DOI:** 10.64898/2026.07.10.737869

**Authors:** Yuanwang Zhang, Hongming Li, Yong Fan

## Abstract

Neurodegenerative diseases exhibit substantial heterogeneity, complicating both diagnosis and prognosis. Identifying clinically meaningful subtypes is crucial for understanding disease mechanisms and can also improve diagnostic precision and prognostic accuracy. Existing subtyping approaches primarily rely on unsupervised learning of patient data for capturing inter-individual variability, often failing to uncover subtypes that are informative for diagnosis or prognosis. To address this limitation, we propose a novel mixture-of-experts (MoE) framework that integrates predictive modeling with subtype identification. Unlike traditional subtyping methods, our approach learns a router to assign individuals to specialized expert networks, each corresponding to a distinct subtype, to improve predictive accuracy. This MoE framework ensures that the discovered subtypes are not only statistically distinct but also clinically informative. We evaluate the framework on a real-world dataset of mild cognitive impairment (MCI) subjects and a semi-simulated dataset, demonstrating superior performance for predicting MCI subjects’ progression to Alzheimer’s disease while identifying distinct clinically meaningful MCI subtypes. Code is available at https://github.com/Kateridge/MoESubtyping.

## 1 Introduction

Neurodegenerative diseases, such as Alzheimer’s disease, are complex disorders characterized by long prodromal phases and progressive deterioration. Disease progression is typically associated with changes in various biomarkers, including volumetric measurements from brain imaging and behavioral or cognitive measures from psychometric assessments [2]. These biomarkers have been extensively studied for their diagnostic and prognostic value [1]. However, neurodegenerative diseases are biologically heterogeneous, and affected individuals may exhibit distinct patterns in in-vivo biomarkers [19,13], posing significant challenges for their effective use in disease diagnosis and prognosis of disease progression. As shown in Fig. 1(a), most existing methods use an all-in-one model to make predictions from heterogeneous data, which may lead to suboptimal prediction performance. Identifying disease subtypes is therefore essential to enhance understanding of the diseases and improve the effectiveness of these data to guide targeted interventions in the prodromal stages.

**Fig. 1.**
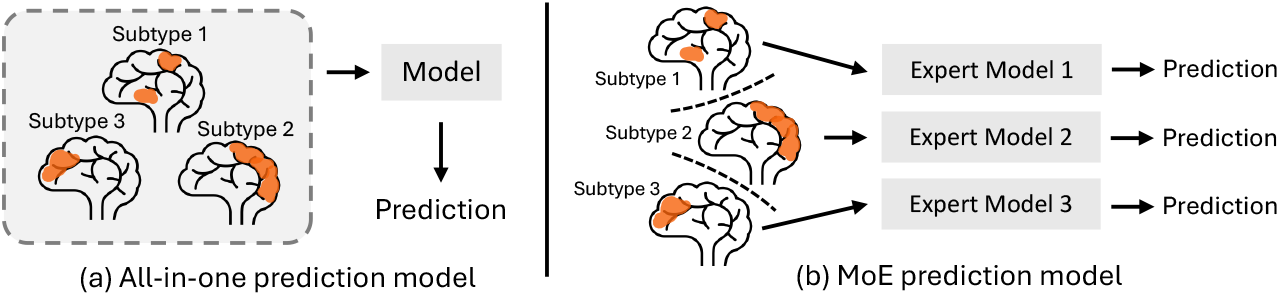
(a) Most existing studies use a single all-in-one model to make predictions from heterogeneous data. (b) Our proposed MoE framework clusters heterogeneous data, routes samples to specialized expert models, which improves the prediction performance while providing interpretable insights into data heterogeneity.

To identify disease subtypes, many studies have applied unsupervised clustering methods, such as K-Means and Gaussian Mixture Models (GMMs), to group individuals based on similarity of measurements derived from brain imaging data [22,7,26]. However, variability in imaging measures does not necessarily correlate with diagnosis or prognosis, meaning that these clusters often capture inter-subject differences without providing informative insights for these downstream tasks. Several studies have also proposed semi-supervised approaches [24,25], which cluster diseased individuals relative to a reference group, such as cognitively unimpaired (CU) subjects, using generative modeling techniques. While these methods achieve promising clustering performance, they still lack discriminative power for prediction tasks. In other words, they can cluster diseased individuals relative to a control group but do not necessarily effectively distinguish diseased from control individuals. Alternative approaches integrate classification and clustering within a unified framework by employing ensembles of multiple SVM classifiers [11,23]. In these models, each linear hyperplane represents a potential subtype, and their ensemble collectively forms a nonlinear decision boundary to distinguish diseased individuals from the reference group. Although these methods enable simultaneous subtype identification and classification, the clustering remains primarily driven by feature similarity rather than predictive relevance.

Mixture-of-experts (MoE) architectures have recently emerged as a promising approach in large language models to scale up the model capacity with fewer computational cost [4]. Many works indicate the MoE model inherently learns the underlying clustering structure of the input distribution driven by the prediction objective [20,17,9]. Inspired by these works, we develop a MoE framework that jointly performs prediction and clustering for heterogeneous neuroimaging data, as illustrated in Fig. 1(b). Unlike prior methods, our approach leverages predictive objectives to guide disease subtype discovery by assigning individuals to specialized expert networks that capture distinct disease patterns. Specifically, as shown in Fig. 2, the framework consists of a router network and multiple expert networks. Each expert performs the prediction task and represents a disease subtype, while the router learns to assign each subject to the most appropriate expert. This design integrates prediction with clustering, enabling more accurate subtype identification while improving overall predictive performance. We extensively validate the proposed framework, demonstrate substantial performance gains on two downstream tasks, and show that these improvements stem from the inherent clustering capability of the MoE architecture. We also derive four distinct patterns of brain atrophy in mild cognitive impairment (MCI) subjects aligned with different disease stages and subtypes.

**Fig. 2.**
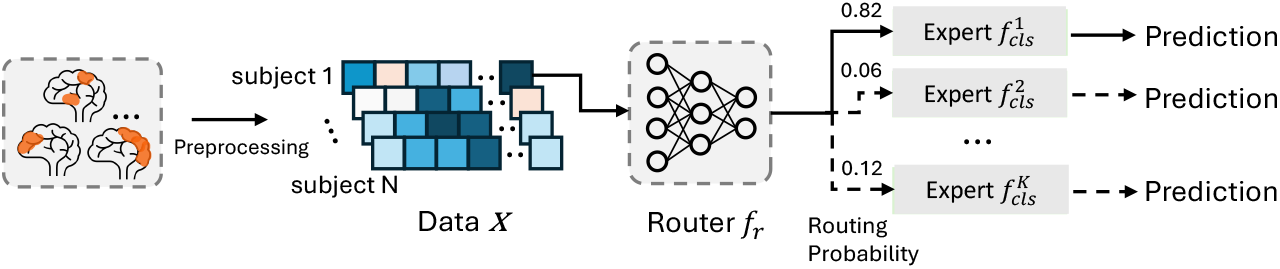
Prediction-driven subtyping using a MoE framework. The MoE architecture assigns individuals to specialized expert networks, each representing a distinct subtype. These expert assignments naturally define cluster memberships, allowing the framework to perform prediction and subtyping simultaneously. This also leads to improved prediction performance compared to all-in-one prediction model.

## 2 Method

In this section, we present the proposed MoE framework, as shown in Fig. 2, along with additional regularization losses designed to improve robustness and prevent trivial solutions.

### 2.1 Mixture-of-experts (MoE) framework

Given N subjects, each represented by a vector of brain regional measurements *X*_*i*_ ∈ ℝ^M^ (*i* = 1, 2, …, N), our MoE model assigns individual subjects to a specialized expert networks 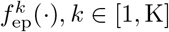, where M is the feature dimension and K is the number of experts. The assignment is determined by a router network *f*_r_(*·*), which outputs an assignment probability vector for each input *X*_*i*_. Each subject is then routed to the expert with the highest probability. These expert assignments naturally define cluster memberships *S*_*k*_ for the input data.

In this study, we implement the MoE framework for two downstream applications: (1) classification of stable MCI versus progressive MCI, and (2) predicting the progression risk of individuals from MCI to dementia. The MoE framework is trained using a task-specific prediction loss, denoted as *L*_task_. For the classification task, each expert is a classifier, and the prediction loss *L*_task_ is defined by the binary cross-entropy loss:

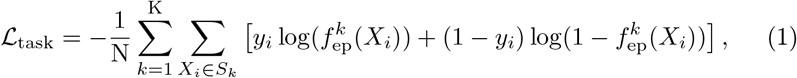

where *y*_*i*_ denotes the binary label of subject *i*, and *S*_*k*_ denote the set of subjects in cluster *k*. For the prediction of conversion risk, each expert is implemented as a Cox regression model, and the loss is defined by the partial likelihood of the Cox proportional hazards model [10]:

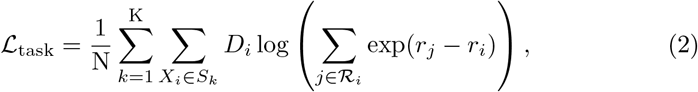

where 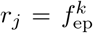 and 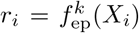 represent the estimated risks of *X*_*j*_ and *X*_*i*_, *D*_*i*_ is the event indicator for subject *i*, and ℛ_*i*_ denotes the set of subjects which are uncensored and have not experienced the event prior to subject *i*.

### 2.2 Regularization losses

Training the model solely with the prediction loss can lead to degenerate router networks. To mitigate this, we introduce three regularization loss functions.

#### Load balance

The MoE framework often suffers from expert collapse, where most or all subjects are routed to a single expert. To prevent expert collapse, we adopt a load-balancing loss [8] that promotes balanced utilization of experts:

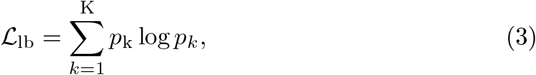

where 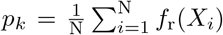 is the average assignment probability for expert k. Minimizing the negative entropy of *p*_*k*_ across experts encourages the router to distribute subjects uniformly among experts, preventing expert collapse.

#### Sparse assignments

Another degenerate case occurs when the router network produces similar assignment probabilities across all experts, resulting in ambiguous routing decisions. To prevent this, we adopt a sparse assignment loss, which encourages each subject to be strongly associated with a single expert. This is achieved by minimizing the entropy of the assignment probabilities over all subjects:

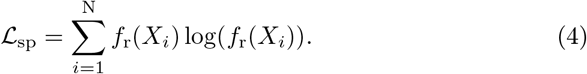

#### Clustering guidance

At the start of optimization, both the router and expert networks are randomly initialized, making it difficult for the router to learn meaningful patterns based solely on the prediction loss. To facilitate convergence and avoid degenerated situations, we introduce a clustering guidance loss *L*_guide_, which uses K-Means assignments as pseudo-labels to guide the router during early training. This loss is defined as the cross-entropy between the router’s routing probabilities and the K-means cluster assignments. The loss weight is gradually decayed to zero over the first 20 epochs.

The overall objective is the weighted sum of all loss items: *L* = *λ*_1_ *L*_task_ + *λ*_2_ *L*_lb_ + *λ*_3_ *L*_sp_ + *λ*_4_ *L*_guide_, where *λ*_1_, *λ*_2_, *λ*_3_ and *λ*_4_ are hyperparameters used to control the strength of losses.

## 3 Experiments

### 3.1 Experiments setup

#### Datasets and preprocessing

We collected 3,458 subjects with 11,602 longitudinal sessions from three cohorts: the Alzheimer’s Disease Neuroimaging Initiative [21], the Open Access Series of Imaging Studies [18], and the Australian Imaging Biomarkers and Lifestyle Study of Aging [12]. Each session included valid diagnostic information, with 5,937 sessions labeled as cognitively unimpaired (CU), 4,703 as mild cognitive impairment (MCI), and 962 as dementia.

To construct the classification dataset, we selected 668 subjects with baseline MRI scans. All selected subjects were diagnosed with MCI at baseline and were further categorized as stable MCI (no progression to dementia with a follow-up period longer than three years) or progressive MCI (progress to dementia within three years). To construct the progression risk prediction dataset, we selected 1,243 subjects with baseline MRI scans and follow-up diagnostic records. Each subject had a baseline diagnosis of MCI and at least two follow-up sessions.

Raw T1-weighted MR images were preprocessed using the standard FreeSurfer^3^ pipeline to extract cortical thickness measures from 68 cortical regions based on the Desikan–Killiany atlas, as well as volumes from 28 subcortical regions based on the Aseg atlas.

#### Semi-simulated dataset

Since ground-truth subtype labels are unavailable in real-world datasets, we further evaluated clustering performance using a semi-simulated dataset. This dataset was generated by imposing simulated brain atrophy patterns on real CU data. Specifically, all CU sessions were randomly divided into a reference group of 2,937 sessions and a simulated disease group of 3,000 sessions. Within the simulated disease group, we created three subtypes, each containing 1,000 sessions, by manually introducing distinct atrophy patterns. For the first subtype, 15 cortical ROIs were randomly selected; for the second subtype, 5 subcortical ROIs were randomly selected; and for the third subtype, 15 cortical and 5 subcortical ROIs were randomly selected. The values of the selected ROIs were then reduced by 10–30%, 10–20%, or 5–15% across different experimental settings, with the atrophy rate for each subject uniformly sampled within the specified range.

#### Model architecture and experimental setting

Each expert and the router network were implemented as a three-layer MLP with an input dimension of 96 and a hidden dimension of 128. ReLU activation and dropout regularization were applied between layers. All models were trained using the SGD optimizer. Experimental results were obtained through 10-fold cross-validation to ensure robust performance evaluation. For the classification task, accuracy (ACC) and sensitivity (SEN) were reported. For the progression risk prediction task, the concordance index (C-index) and the averaged time-dependent AUC [3] were used as evaluation metrics. For clustering performance on the semi-simulated dataset, the adjusted Rand index (ARI) and normalized mutual information (NMI) were reported.

### 3.2 MoE improves prediction performance

#### Comparison to baselines

The baseline all-in-one prediction model (Single) was implemented using the same MLP architecture with a single expert network. To ensure a fair comparison with the MoE model (which has a larger number of parameters), we also implemented a higher-capacity baseline in a similar model structure with 5× parameters (Single*) by increasing the size of the hidden layers. As shown in Table 1, the MoE framework substantially outperforms both the standard and large all-in-one models on both tasks. These results highlight the benefits of clustering inputs and training specialized expert networks.

**Table 1.** Prediction performance for classification and progression risk prediction. Mean metrics across folds with 95% confidence interval were reported.

| Methods | Classification |  | Progression Risk Prediction |  |
| --- | --- | --- | --- | --- |
| | Accuracy ↑ | Sensitivity ↑ | C-Index ↑ | AUC ↑ |
| Single | 0.729 (0.694-0.763) | 0.710 (0.649-0.770) | 0.725 (0.696-0.753) | 0.787 (0.754-0.821) |
| Single* | 0.736 (0.714-0.759) | 0.699 (0.649-0.750) | 0.738 (0.712-0.764) | 0.803 (0.740-0.832) |
| MoE (2 experts) | 0.750 (0.716-0.783) | 0.720 (0.667-0.774) | 0.750 (0.722-0.779) | 0.813 (0.784-0.842) |
| MoE (3 experts) | **0.769 (0.741-0.797)** | 0.726 (0.676-0.776) | 0.757 (0.733-0.781) | 0.817 (0.792-0.843) |
| MoE (4 experts) | 0.765 (0.737-0.794) | **0.761 (0.699-0.823)** | **0.763 (0.740-0.786)** | **0.830 (0.802-0.858)** |
| MoE (5 experts) | 0.761 (0.736-0.786) | 0.737 (0.675-0.800) | 0.756 (0.730-0.782) | 0.819 (0.790-0.847) |
| Random Forests | 0.732 (0.700-0.763) | 0.671 (0.607-0.734) | 0.749 (0.727-0.770) | 0.814 (0.789-0.839) |
| TabPFN [14] | 0.768 (0.732-0.803) | 0.722 (0.659-0.786) | - | - |
| DeepSurv [16] | - | - | 0.729 (0.707-0.752) | 0.796 (0.764-0.828) |

#### Effects of cluster number

We further investigated the impact of the number of experts/clusters, as shown in Table 1. Performance improved as the number of experts increased and peaked at 4 experts, suggesting that these MCI subjects may naturally group into 4 distinct subtypes.

#### Compare to other methods

We further compared our approach with several established methods: (1) Random Forests for both tasks, (2) a tabular foundation model, TabPFN [14], for the classification task, and (3) a widely used survival analysis model, DeepSurv [16], for the progression risk prediction task. As shown in Table 1, our method outperformed Random Forests and DeepSurv, while achieving performance comparable to TabPFN.

### 3.3 MoE inherently characterizes heterogeneous patterns of MRI

We further demonstrate that the performance improvements of the MoE model can be attributed to its inherent ability to cluster inputs.

#### Clustering performance on semi-simulated dataset

The clustering capability was evaluated using the semi-simulated dataset. The prediction task was formulated as binary classification between the reference group and the simulated disease group. To ensure a robust evaluation, we used clustering results from different methods as target labels of the clustering guidance loss, including K-means, Gaussian mixture models (GMM), and DecoNet [15]. As shown in Table 2, the MoE framework consistently improves clustering performance across different atrophy rate settings. These results indicate that the MoE model can inherently cluster inputs.

**Table 2.** Clustering performance on the semi-simulated dataset. Results are reported as the agreement between predicted subtypes and the ground-truth subtype labels.

| Atrophy Rate | 10%-30% |  | 10%-20% |  | 5%-15% |  |
| --- | --- | --- | --- | --- | --- | --- |
| | ARI ↑ | NMI ↑ | ARI ↑ | NMI ↑ | ARI ↑ | NMI ↑ |
| K-Means | 0.937 | 0.916 | 0.640 | 0.636 | 0.289 | 0.367 |
| MoE+K-Means | **0.998** | **0.997** | **0.717** | **0.719** | **0.419** | **0.476** |
| GMM | 0.998 | 0.997 | 0.677 | 0.692 | 0.408 | 0.478 |
| MoE+GMM | **1.000** | **1.000** | **0.713** | **0.720** | **0.451** | **0.504** |
| DecoNet [15] | 0.998 | 0.997 | 0.647 | 0.674 | 0.513 | 0.579 |
| MoE+DecoNet | **0.999** | **0.997** | **0.656** | **0.688** | **0.529** | **0.592** |

#### Ablation study for regularization losses

We also evaluated the impact of each regularization loss on the MoE model, as summarized in Table 3. Without any regularization, the MoE model can converge to degenerate solutions that fail to cluster inputs, and its prediction performance collapses to that of a standard all-in-one model. Introducing the regularization losses improves both clustering and prediction performance. These results demonstrate the effectiveness of each loss component and support the conclusion that the MoE’s predictive gains are closely linked to its inherent clustering ability.

**Table 3.** Ablation study demonstrating the contributions of different regularization losses. Clustering performance was evaluated on the semi-simulated dataset with 10%–20% atrophy rates, and prediction performance was assessed on both the classification and progression risk prediction tasks using 4 experts.

| Regularization | Clustering (ARI) | Clustering (NMI) | Classification (ACC) | Progression Risk Prediction (C-Index) |
| --- | --- | --- | --- | --- |
| $\mathcal{L}_{\text{task}}$ only | 0.000 | 0.000 | 0.732 | 0.745 |
| $\mathcal{L}_{\text{task}} + \mathcal{L}_{\text{lb}}$ | 0.145 | 0.129 | 0.739 | 0.750 |
| $\mathcal{L}_{\text{task}} + \mathcal{L}_{\text{lb}} + \mathcal{L}_{\text{sp}}$ | 0.312 | 0.307 | 0.751 | 0.753 |
| $\mathcal{L}_{\text{task}} + \mathcal{L}_{\text{lb}} + \mathcal{L}_{\text{sp}} + \mathcal{L}_{\text{guide}}$ | **0.717** | **0.719** | **0.765** | **0.763** |

### 3.4 Four subtypes identified in MCI population

Finally, we set the number of experts to 4 for the progression risk prediction task and grouped MCI subjects into 4 subtypes based on assignments from the router. We then computed the effect size for each ROI relative to the CU population and visualized the resulting patterns on cortical Desikan–Killiany and subcortical Aseg brain atlases, as shown in Fig. 3. Subtype 4 exhibits severe atrophy across both cortical and subcortical regions, consistent with dementia-related pathology (late-stage MCI), whereas Subtype 1 shows only mild atrophy, consistent with early-stage MCI. Subtype 3 demonstrates widespread cortical atrophy, while Subtype 2 presents prominent atrophy in the medial temporal lobe, consistent with an amnestic MCI pattern.

**Fig. 3.**
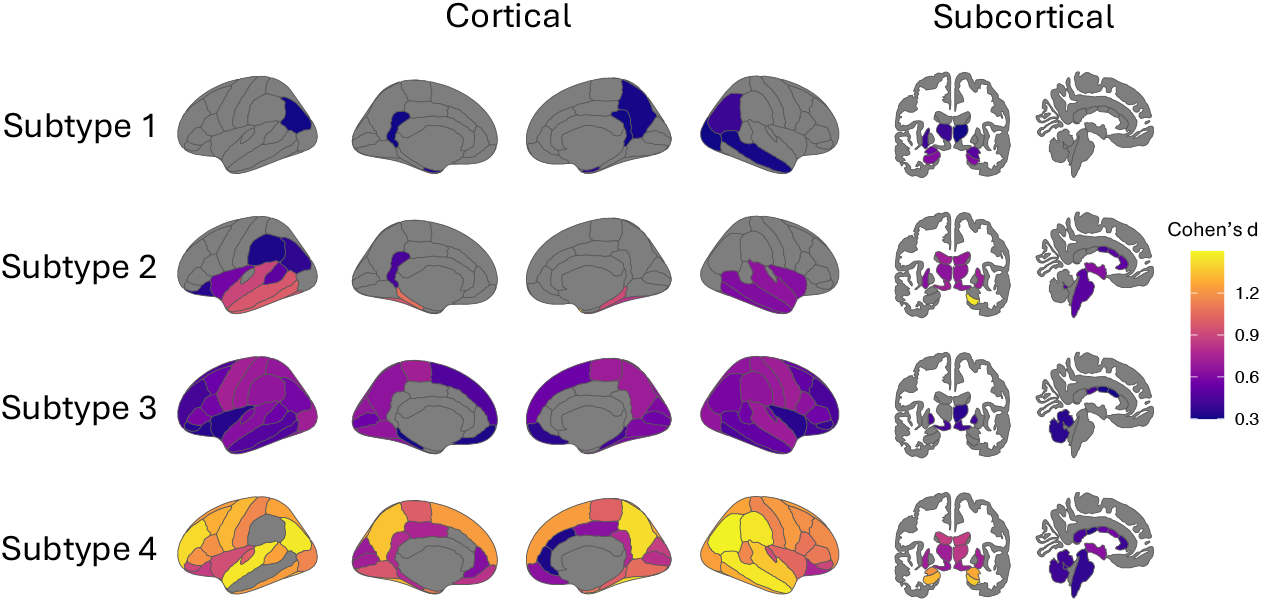
Cortical and subcortical visualization of effect sizes (Cohen’s d) for different MCI subtypes relative to the CU population. Higher values indicate greater differences compared to CU subjects.

## 4 Discussion and Conclusions

We developed a generalized MoE framework for joint prediction and subtype identification in neurodegenerative diseases. By leveraging multiple expert networks, the framework simultaneously enhances predictive performance and un-covers meaningful subtypes. Furthermore, the identified subtypes revealed distinct patterns of brain atrophy aligned with different disease stages and subtypes, underscoring the potential of prediction-driven subtyping for advancing precision medicine in neurodegenerative disorders.

Our study could be further improved in future work to enhance generalizability and applicability. First, our framework is readily applicable to voxel-wise imaging data, by replacing the expert networks with image-based backbones and substituting the clustering guidance model with other deep learning–based clustering methods [6,5]. This can potentially improve the performance and generalization as voxel-wise images may contain richer features. Second, the clustering guidance loss was primarily designed to provide a good initialization for the router, improving convergence and preventing degenerate solutions. Future work could explore incorporating fully optimizable clustering objectives directly into the framework, which may further enhance performance if the prediction and clustering objectives are appropriately balanced.

## Footnotes

3 https://surfer.nmr.mgh.harvard.edu/

